# Pleistocene Mammal Population Fluctuation Patterns Inferred by Their Genomes

**DOI:** 10.1101/505131

**Authors:** Yulu Liu, Biao Liu, Xingxin Pan, Qiong Shi, Zhoujian Xiao, Shengbin Li, Shuaicheng Li

## Abstract

Paleoclimate fluctuations critically affect paleoecological systems and influence mammal populations, even resulting in population differentiation [1]. Historical effective population size (*N*_*e*_) can reflect these influences [2, 3]. Dozens of recent studies have investigated the relationship between variations in *N*_*e*_ values of one or a small number of mammalian species, inferred from genomic data, and fluctuations in paleoclimate [4-7]. However, there lacks an integrated and comprehensive study on the relationship between the fluctuations in paleoclimate and variations in *N*_*e*_ values inferred from genome sequencing data of a wide range of mammals. To investigate patterns in mammalian *N*_*e*_ values during the the Pleistocene, we gathered whole genome sequencing data of 60 mammals from 35 species distributed across Afro-Eurasia and the Americas, then inferred their *N*_*e*_ curves using the Pairwise Sequentially Markovian Coalescent (PSMC) method; 30 mammalian *N*_*e*_ curves almost simultaneously started to contract at the turning point of the Middle Pleistocene Transition (MPT); then the population of seven mammals started to expand at the turning point of the Middle Brunhes Event (MBE), while the contraction of other mammals’ populations was prolonged to the later different time periods. Eight mammals experienced a severe population contraction around the Last Glaciation Maximum, as some aves did [8], while four potential ruminant beneficiaries showed an expanding population. *Sus scrofa* and *Bos taurus* experienced an internal population differentiation in the MPT. To conclude, the phenomenon that critical paleoclimate events facilitated contemporaneous animal population fluctuations in the paleoecological system is showed by our *N*_*e*_ curve analysis.

## RESULTS AND DISCUSSION

### Effective population size of mammals during the Pleistocene

By utilizing the nonuniform distribution of single nucleotide variants (SNVs) in the genome, the Pairwise Sequentially Markovian Coalescent (PSMC) method can reconstruct the historical effective population size (*N*_*e*_) from any single individual of the population [4]. In this work, we applied the PSMC method to obtain the *N*_*e*_ curves of 35 mammal species in Afro-Eurasia and the Americas (Figure S1). The *N*_*e*_ curves estimated from the genome data span the whole Pleistocene, yielding us that were sufficient to conduct the subsequent analysis.

### Climates contributed to the mammal N_e_ curves

*N*_*e*_ fluctuation could result from a combined effect of genetics, climate, locations, or other factors [8]. To understand the degree that climate contributed to the *N*_*e*_ curves, we extracted the principal components (PCs) from 60 *N*_*e*_ curves by the principal component analysis (PCA). PC1 and PC2 contributed variance ratios of 0.508 and 0.273, respectively (Figure 1A). PC1 accounts for most of the variance; however, after hierarchical clustering, no signals could be detected to support that PC1 is correlated with genetic background (Figure S2). This finding could suggest that the complex paleoclimate impact on the mammals. We clustered these mammals hierarchically with PC2 (Figure 1B) and discovered that in many mammals (*Ursus, Ponginae, Homo, Canis*, some subgroups in *Rhinopithecus* and some herbivorous mammals), their genetically close subgroups were clustered in the same branch, with the exception of *Bos* and *Sus*. Eurasian and African *Bos* clusters as well as Asian and European *Sus* clusters are separated by a clear boundary, reflecting a long separation time of their populations that will be elaborated in the following section. Hence, PC2 could be interpreted as the integrated effect of the inherent genetic characteristics and geographical locations on the mammals. In general, close locations imply similar climate conditions; therefore, fluctuations of mammal *N*_*e*_ curves can reflect the climates of the mammal ancestors confronting to some extent.

**Figure 1.**
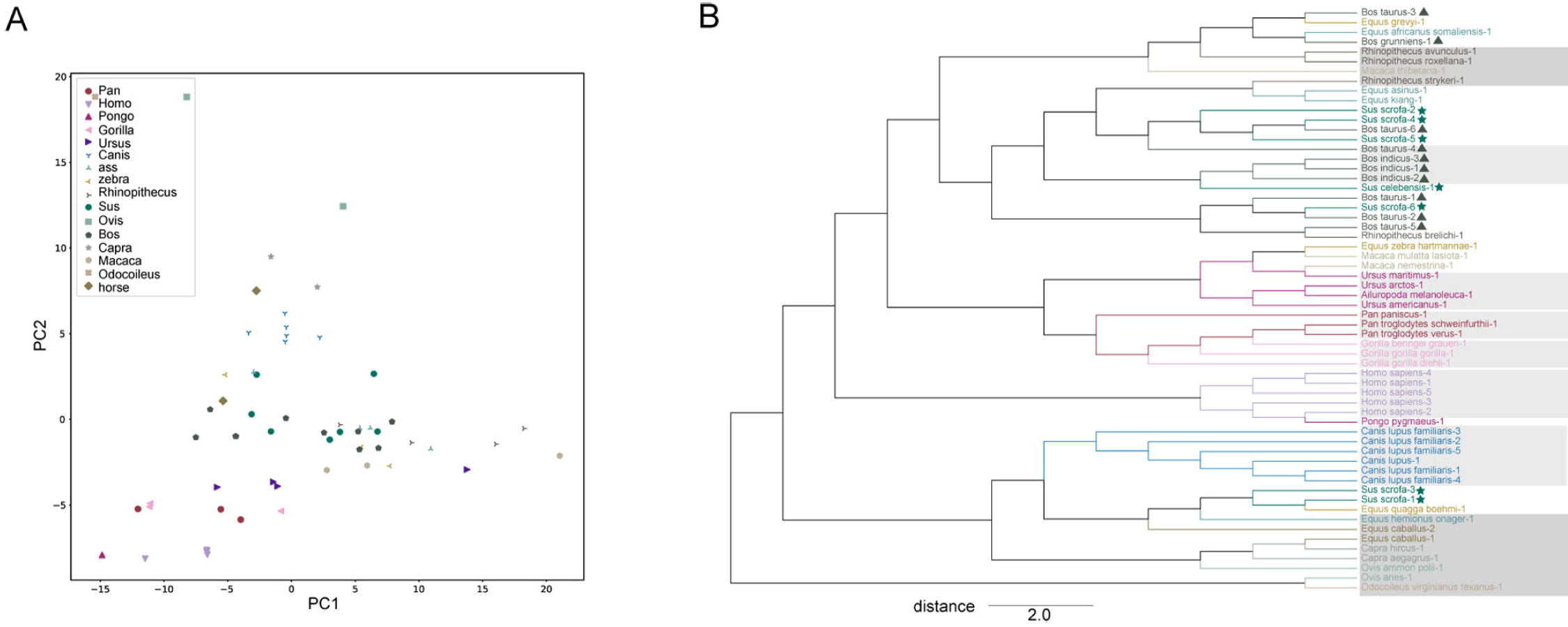
PC analysis of mammal demographic trajectories. (A) PC1 and PC2 plots. Different colors or shapes represent different mammal classes. (B) Clusters obtained using PC2 of *N*_*e*_ curves. Colors represent mammal classes. *Bos* and *Sus* are marked by triangles and pentagrams, respectively. Mammals in light gray shading share a close genetic relationship and cluster closely when using PC2 of *N*_*e*_ curves. Mammals in dark gray shadows share a relatively close genetic relationship, live in similar habitats, and cluster closely when using PC2 of *N*_*e*_ curves. The scale bar represents the Euclidean distance between samples.

**Figure 2.**
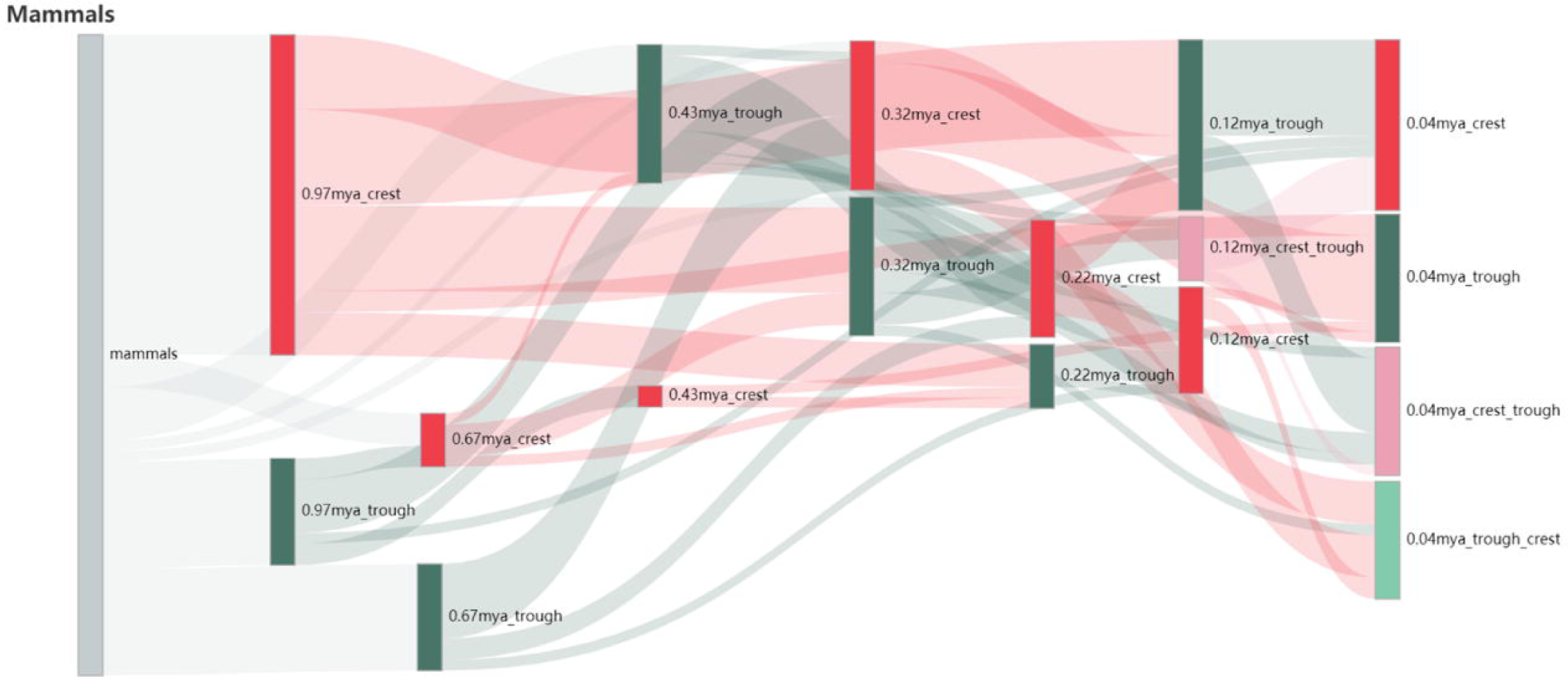
Sankey diagram of mammal extrema. Mammals in the same column possess the same extremum. Sample names are in Table S1.

### N_e_ curve extrema clustered into seven clusters

*N*_*e*_ curve extrema are consistent with significant climate events in the Pleistocene. These extrema (Table S1) are centered at seven time points with an average silhouette coefficient of 0.64, --an acceptable cluster indicator. All seven clusters coincide with turning points of important climate events, as elaborated below.

The earliest cluster centered at 0.97±0.021 mya, with 40 extrema, coincided with the turning point of the Middle Pleistocene Transition (MPT, 0.94 mya [9]). We refer to this cluster as the MPT Cluster. In the MPT Cluster, 30 *N*_*e*_ curves reached a peak, including those of *Sus, Bos, Ovis, Capra, Equus caballus, Ailuropoda melanoleuca, Rhinopithecus roxellana, Rhinopithecus strykeri, Ursus arctos, Ursus maritimus, Pan troglodytes verus* and *Gorilla gorilla diehli*; and ten *N*_*e*_ curves displayed a bottleneck, including *Homo* and five species from *Equus*. Accordingly, we categorized the mammals into three classes (Table S1): MPT Maxima Class, MPT Minima Class, and MPT Neutral Class (mammals whose *N*_*e*_ curves are without the first extremum in the MPT class).

The contraction of the MPT Maxima Class around 0.97±0.021 mya may result from the Jaramillo geomagnetic reversal (1.07 mya∼0.99 mya) [10] or changes in solar insolation after the coeval d13C_max-III_ event of 1.0∼0.95 mya (Marine isotope stage (MIS) 29–25) [11, 12]. The climate became too harsh at a large scale for mammals to survive [9] and led to massive mammal extinction or population contraction events such as warm-steppic mammals being replaced with cold-steppic mammals in European Villafrancian fauna [13] and the southern-migration mammal wreck event of Gongwangling fauna [14]. The Plio-Pleistocene contributed substantially to current biomes [15] and MP fossils of 73.3% mammals in this class, including *Ailuropoda* [16], *Bos* [17], *Capra* [18], *Equus* [19], *Rhinopithecus* [20] and *Sus* [21] are majorly distributed in the Palearctic, according to the fossil database (Figure S3A, B, D, E, K and L). Therefore, the population peak of the MPT Maxima Class in the MPT Cluster suggests that the ancient populations of MP mammals in the Palearctic could have experienced severe contractions around the turning point of the MPT.

The MPT Maxima Class population started to contract from the MPT Cluster center, and their first bottlenecks after that are distributed in five clusters (Figure 3A). The MPT Maxima Class, including African *Pan troglodytes verus* and *Bos*, wild-spread *Equus* and European *Sus Scrofa,* reached the first bottleneck at 0.43±0.018 mya, which is referred to as the MBE (Middle Brunhes Event) Bottleneck for 0.43 mya is happens to be the turning point of the MBE (MIS13-MIS11) [22]. Eight mammals, including *Capara*, Asian *Sus scrofa, Ovis aries*, and *Bos indicus*, reached the first bottleneck of their population at approximately 0.32±0.01 mya, the time which was known during the Mindel-Riss interglacial [23] (MRI bottleneck). These mammals, except the *Bos indicus*, were sampled at a relatively high latitude. In the Mindel-Riss interglacial, the climate was similar to the current climate. For four mammals including *Ursus arctos, Ursus maritimus* and two *Bos taurus* individuals, their ancestral *N*_*e*_ values synchronously reached the first bottleneck at 0.22±0.010 mya in the middle of the Riss glaciation, which is a relatively weak MIS 7 period with a cool interglaciation climate [24] (MRG Bottleneck). For nine mammals from the MPT Maxima Class, including *Ovis ammon polii, Ailuropoda melanoleuca, Bos grunniens, Rhinopithecus roxellana* and *Sus celebensis* as well as three *Bos taurus* individuals and one *Sus scrofa* individual, their ancestral *N*_*e*_ values started to show an expansion of their populations until 0.12±0.008 mya in the warmest, last interglaciation, the Eemian interglaciation [25] (EI Bottleneck). Two species, *Gorilla gorilla diehli* and *Rhinopithecus strykeri*, reached the first bottleneck until 0.04±0.004 mya (the most recent cluster center). These mammal populations contracted almost simultaneously but started to expand at different time period.

**Figure 3.**
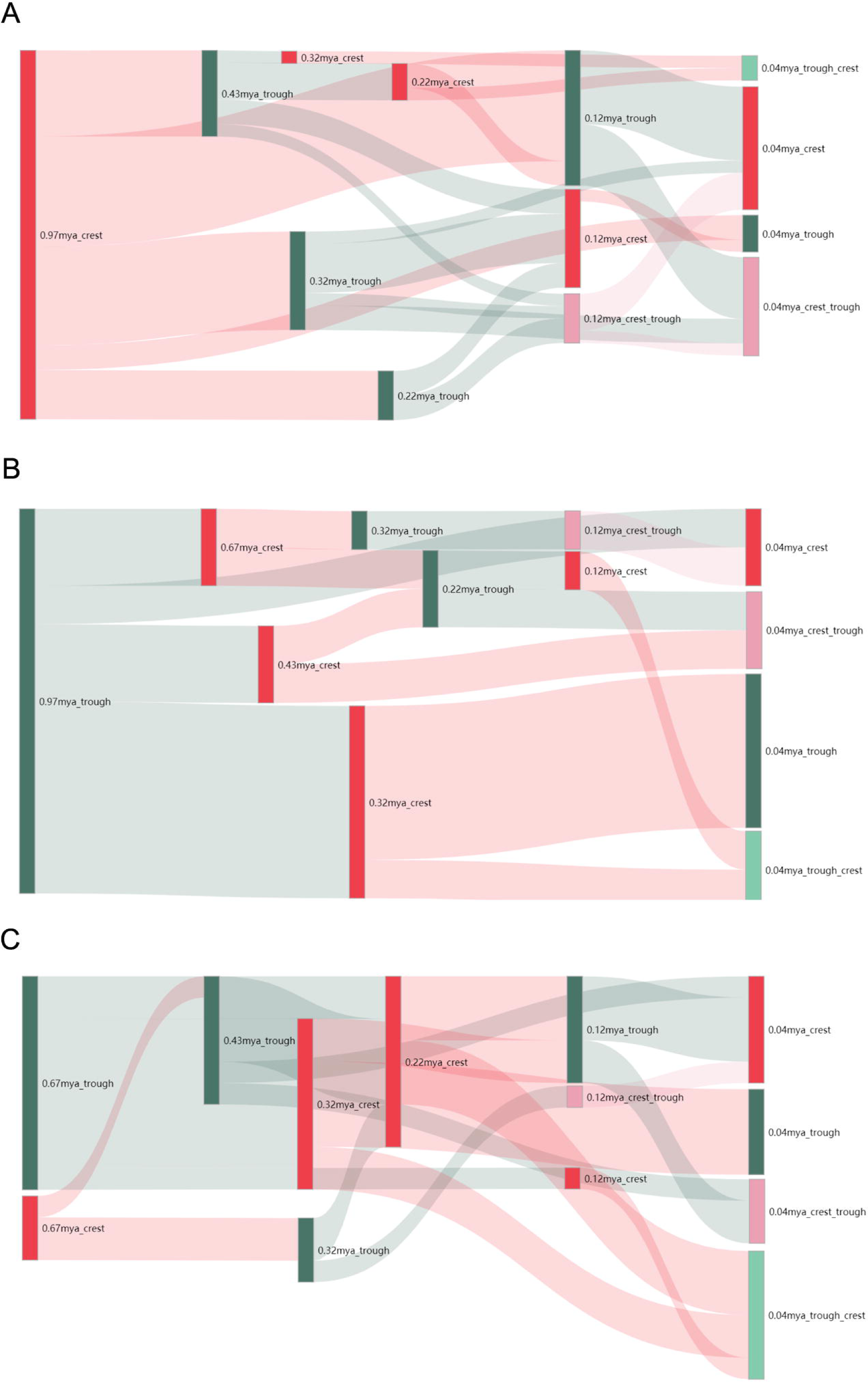
Sankey diagram of three classes. (A) MPT Maxima Class. (B) MPT Minima Class. (C) MPT Neutral Class. Sample names are also in Table S1.

Climate conditions could prompt mammal population migration and differentiation. As a result, the population of migrated mammals may have started to expand, while mammals that did not migrate sustained contraction. In the MPT Maxima Class, the mammals that experienced the MBE Bottleneck likely lived in low latitude regions, and those that experienced the MRI Bottleneck usually resided at relatively high latitudes with the exception of *Bos* and *Sus scrofa*, which will be elaborated in the following section.

However, there was no obvious association between population fluctuations and habitats of the mammals that experienced the MGR and EI bottlenecks. The fact that mammals in the MPT Maxima Class spread in different latitudes, adapted themselves to different environments and developed different habits, but shared five asynchronous expansion times, could result from combined factors of environment in different latitudes and the capacity for mammalian adaptations.

We also noticed that ten mammals of the MPT Neutral Class (Figure 3C) reached a bottleneck while three of them reached a peak, at 0.67±0.030 mya, or the end of the MPT. At that time, the planet held a high ice volume but was relatively warm [24, 27]. The mammals that underwent a bottleneck around 0.67±0.030 mya included *Pongo pygmaeus, Ursus americanus Macaca mulatta lasiota, Macaca thibetana* and *Canis*. Their populations began to shrink separately before 1 mya, indicating that the population contraction impetus differed for mammals from the MPT Maxima Class. The mammals with a peak at that time included *Odocoileus virginianus texanus, Gorilla gorilla gorilla* and *Gorilla beringei graueri*.

## Differentiation in mammal populations

*Sus scrofa* and *Bos* populations experienced internal population differentiation between 0.97 mya and 0.43 mya in the middle Pleistocene transmission.

*Sus scrofa* belongs to the MPT Maxima Class (Table S1); thereafter, the population of European *Sus scrofa* experienced a prolonged contraction until the MBE Bottleneck, while the population of Asian *Sus scrofa* continued contracting until the MRG Bottleneck (Figure 4A). The middle Pleistocene fossils of Asian *Sus scrofa* were only distributed in relatively low latitude areas in Asia (Figure S3L). We also noticed that some Asian mammals, such as *Capra aegagrus, Capra hircus* [18], and *Ailuropoda melanoleuca* [16], also started their population expansion at the MRG Bottleneck, while the European *Sus scrofa* population started to expand at the turning point of the MBE, which is the same time as the *Equus caballus* population did [19]. The long-term population contraction of Eurasian *Sus scrofa*, following a different fluctuation trend in the MBE Bottleneck, indicates a differentiation in Eurasian *Sus scrofa* during the MPT and MBE. Kijas estimated that Eurasian *Sus scrofa* maternity divergence time is ∼0.9 mya according to the mtDNA genome sequence [32]; with synonymous and nonsynonymous nucleotide substitutions of two main domains of the D-loop region in the cytochrome B gene (CytB) (1,847 bp), Alves estimated the Eurasian *Sus scrofa* divergence time to be ∼0.6 mya [33]. Based on interpopulational distances for the mtDNA CytB sequence (1,140 bp), another estimated divergence time is ∼0.5 mya [33, 34]. With phylogenetic analysis based on pairwise genetic distance of the mitotypes of Eurasian *Sus scrofa*, using an evolutionary rate of 2%, their divergence time was estimated as 0.28 mya; according to mtDNA D-loop sequence divergence time analysis, the Eurasian *Sus scrofa* divergence time is ∼58 kya [35]. Our *N*_*e*_ curves analysis support that Eurasian *Sus scrofa* could have differentiated in 0.9∼0.5 mya (during the long, harsh MPT [9]). The estimations that the divergence time of Eurasian *Sus scrofa* is ∼0.28 mya or ∼58 kya could have resulted from the inadequate sequence lengths used for the respective estimations.

**Figure 4.**
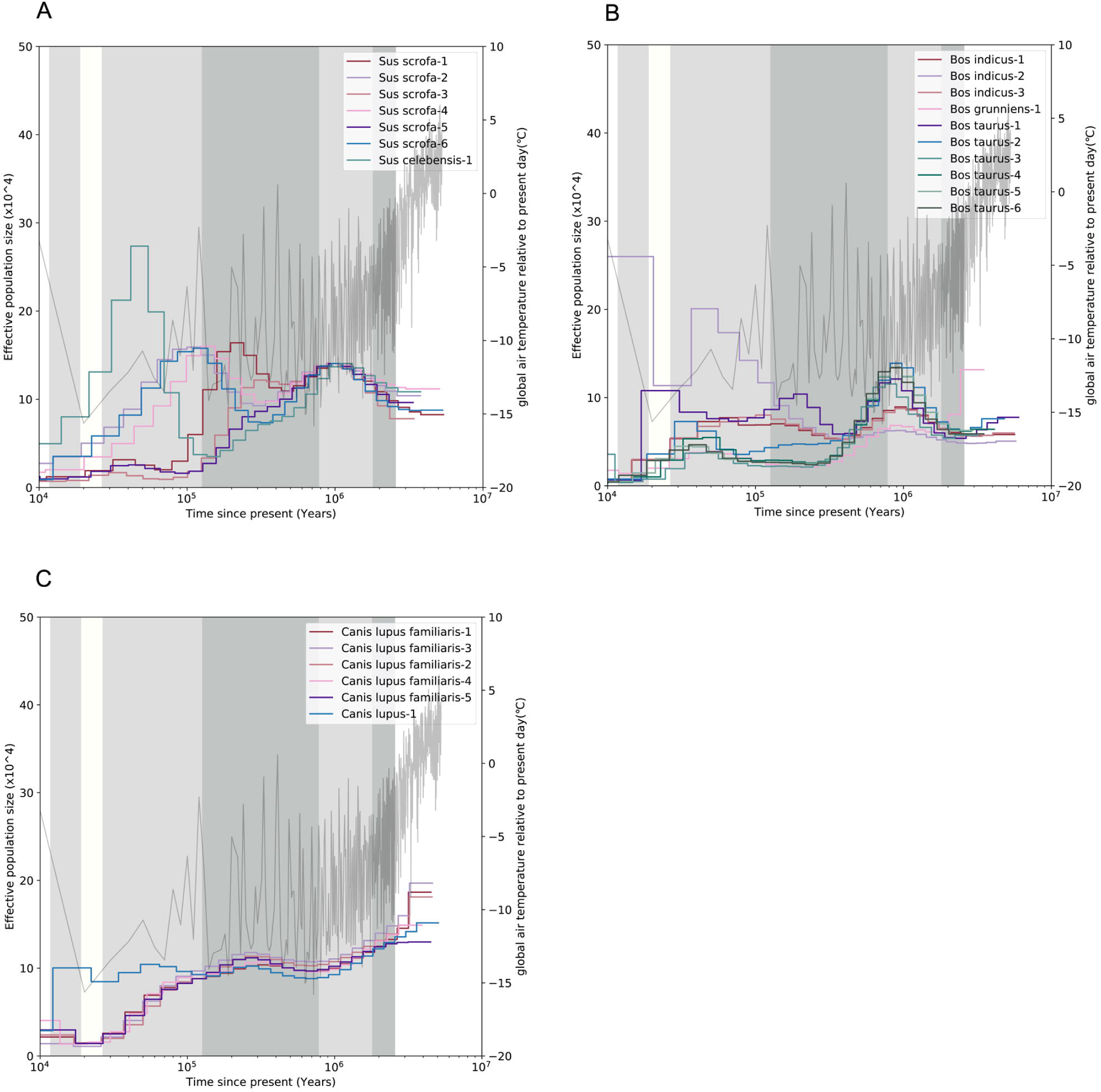
three mammal *Ne* curves. (A)*N*_*e*_ curves of *Sus scrofa* and *Sus celebensis*; (B)*N*_*e*_ curves of *Bos taurus, Bos indicus* and *Bos grunniens*; (C) *N*_*e*_ curves of *Canis lupus familiaris* and *Canis lupus*-1.

We noticed that for all the *Bos* individuals, their ancestral populations peaked at the MPT Maxima Class as *Sus* did, while they experienced their subsequent bottlenecks at different clusters (Figure 3A). The first population bottleneck of African *Bos indicus* and one African *Bos taurus* located in the MBE bottleneck while Eurasian *Bos taurus* reached the first bottleneck at later different cluster (Figure 4B), this phenomenon indicates that the differentiation of the *Bos taurus* population occurred prior to the MBE bottleneck. Since 92% mammals from MPT Maxima class are Palearctic mammals or Indomalayan mammals exclude African *Bos*, the potential *Bos* population could originate in Palearctic. The first *Bos* fossil in Ethiopia originated in ∼0.6 mya, indicating a migration of *Bos* [36]. which is also consistent with our estimated *Bos taurus* differentiation event, and because of their potential ancestral population originated in the Palearctic, we could infer that their migration direction is from Eurasia to Africa. Consequently, the harsh MPT period could also result in the migration and differentiation of *Bos taurus*.

### The fluctuation of mammal N_e_ in the Late Pleistocene (LP) period

The *N*_*e*_ curves of many mammals fluctuated during the LP period. Around the time of the LGM and Younger Dryas Event (YDE) (12.6∼11.6 kya) [28, 29], the ancestral populations of seven mammal species, including *Equus hemionus onager, Macaca nemestrina, Equus quagga boehmi, Canis lupus, Odocoileus virginianus texanus*, and *Equus caballus*, displayed a sudden contraction (Figure S4A). Furthermore, the populations of four species, including *Capra aegagrus, Gorilla gorilla diehli* and non-African *Homo sapiens*, experienced a sharp increase (Figure S4B).

The aforementioned fluctuation was accompanied by a megafauna extinction during which many large mammals extricated [37-40]. The time of coincidence of the two events indicates that the causes of the megafauna extinction event could also have intensively influenced the ancestors who experienced the aforementioned decline event of the extant mammals. The LGM and the YDE could have contributed to the ancient mammal *N*_*e*_ decline event as well as the megafauna extinction event [37]. Many Aves populations also displayed such fluctuations at the same time, as indicated by their *N*_*e*_ curves [8]. However, neither the LGM nor the YDE was solely sufficient to extinguish megafauna [41], nor could they be the only substantial causes for the *N*_*e*_ decline event. Hence, there could be other factors contributing to these two events.

*Homo sapiens* ancestors could also have contributed to the *N*_*e*_ fluctuations. In our results, the *N*_*e*_ curves of *Canis lupus familiaris* and *Canis lupus* started their divergences ∼0.1 mya, corroborating that the estimated divergence time of the early dog and wolf could be as early as 135 kya [42]. The *N*_*e*_ of the ancestor of *Canis lupus* experienced a sudden, sharp decline event around the YDE, while the ancestor of *Canis lupus familiaris* escaped such a decline (Figure 4C). According to recent research, *Canis lupus familiaris* was domesticated in China in ∼33 kya [43], in Central Asia in ∼12.5 kya or in Europe in ∼15 kya [44]. The domestication events occurred around a population contraction event of *Canis lupus,* the ancestor of which lived without domestication by *Homo sapiens*. Therefore, humans could be related to the expansion of the *Canis lupus familiaris* ancestral population. In the last glaciation, *Homo sapiens* started to spread across the world from Africa with brilliant hunting ability in ∼60 kya [45]. Encountering these clever hunters, megafauna seldom understood how to survive. Many mammal *N*_*e*_ decline events also happened after *Homo sapiens* spread, according to our results (Figure S4A). The *Homo sapiens* spreading and occupancies agree with the time and trace for the extinction of mammoths [37, 46, 47]. This hints that *Homo sapiens* could have played an essential role in modern mammal ancestral *N*_*e*_ contractions in the LP. Furthermore, the sharp expansion of *Bos indicus*-2, *Capra aegagrus* and *Gorilla gorilla diehli* after the LGM could also be attributed to mammalian ancestor population LP contractions and megafauna extinctions. Their population contraction event vacated ecological niches; thereafter, hostile environment adaptive ruminants such as *Bos* and *Capra* could have been the potential beneficiaries during this harsh period, and the domestication of *Homo sapiens* could also have facilitated their population expansion after the YDE [48].

## DISCUSSION

Considering earth as a system, paleoclimate fluctuations affect the rise and fall of the population of earth’s creatures in the paleoecological system. *N*_*e*_ curves inferred by PSMC analysis offer a chance to investigate *N*_*e*_ dynamics in ancient times, and fossil distributions can verify these results. The influences of the MPT, MBE, LGM and other paleoclimate events on the paleoecosystem are obviously reflected in the mammal *N*_*e*_ curves. Furthermore, the analysis of these mammal *N*_*e*_ curves reveals preliminary impacts of climate fluctuations on the biosphere. In addition, the mammal *N*_*e*_ decline events in harsh stages alert us to pay attention to the dangers that could be caused by dramatic climate change in the future.

## ACKNOWLEDGMENTS

We are grateful to Yanlin zhang (McGill) for excellent technical help.; andwe thank Bailiang Jian (BGI) and Yilin Liu (BGI) for improving the manuscript;we also acknowledge BGI-Shenzhen for the computing resources; we thank Fossilworks and the contributors who provided fossil records for us, including the following:

A. Behrensmeyer. Taxonomic occurrences of mammal recorded in the fossilworks. http://fossilworks.org.

A. Hendy. Taxonomic occurrences of mammal recorded in the PaleoDB. http://fossilworks.org.

A. Miller. Taxonomic occurrences of Equus recorded in the PaleoDB. http://fossilworks.org.

A. Turner. Taxonomic occurrences of mammal recorded in the PaleoDB. http://fossilworks.org.

C. Bell. Taxonomic occurrences of mammal recorded in the PaleoDB. http://fossilworks.org.

C. Jaramillo. Taxonomic occurrences of mammal recorded in the PaleoDB. http://fossilworks.org.

D. Croft. Taxonomic occurrences of mammal recorded in the PaleoDB. http://fossilworks.org.

E. Fara. Taxonomic occurrences of B recorded in the PaleoDB. http://fossilworks.org.

E. Vlachos. Taxonomic occurrences of Bos recorded. http://fossilworks.org.

J. Alroy. Taxonomic occurrences of mammal recorded in the Fossilworks. http://fossilworks.org.

J. Bloch. Taxonomic occurrences of mammal recorded. http://fossilworks.org.

J. Marcot. Taxonomic occurrences of mammal recorded. http://fossilworks.org.

M. Carrano. Taxonomic occurrences of mammal recorded in the PaleoDB. http://fossilworks.org.

M. Uhen. Taxonomic occurrences of mammal recorded in the PaleoDB. http://fossilworks.org.

P. Holroyd. Taxonomic occurrences of Equus recorded. http://fossilworks.org.

P. Mannion. Taxonomic occurrences of Equus recorded in the PaleoDB. http://fossilworks.org.

W. Clyde. Taxonomic occurrences of mammal recorded in the PaleoDB. http://fossilworks.org.

W. Kiessling. Taxonomic occurrences of Macaca recorded. http://fossilworks.org.

## AUTHOR CONTRIBUTIONS

S.C.L. supervised the research and revised the manuscript; Y.L.L. collected, analyzed data and prepared the manuscript draft; B.L. participated in the literature review and performed some analysis; X.X.P. collected some datasets; Z.J.X provided assistance; Q.S supervised the research; S.B.L revised the manuscripts.

## DECLARATION OF INTERESTS

The authors declare no competing interests.

## STAR★METHODS

### KEY RESOURCE TABLE

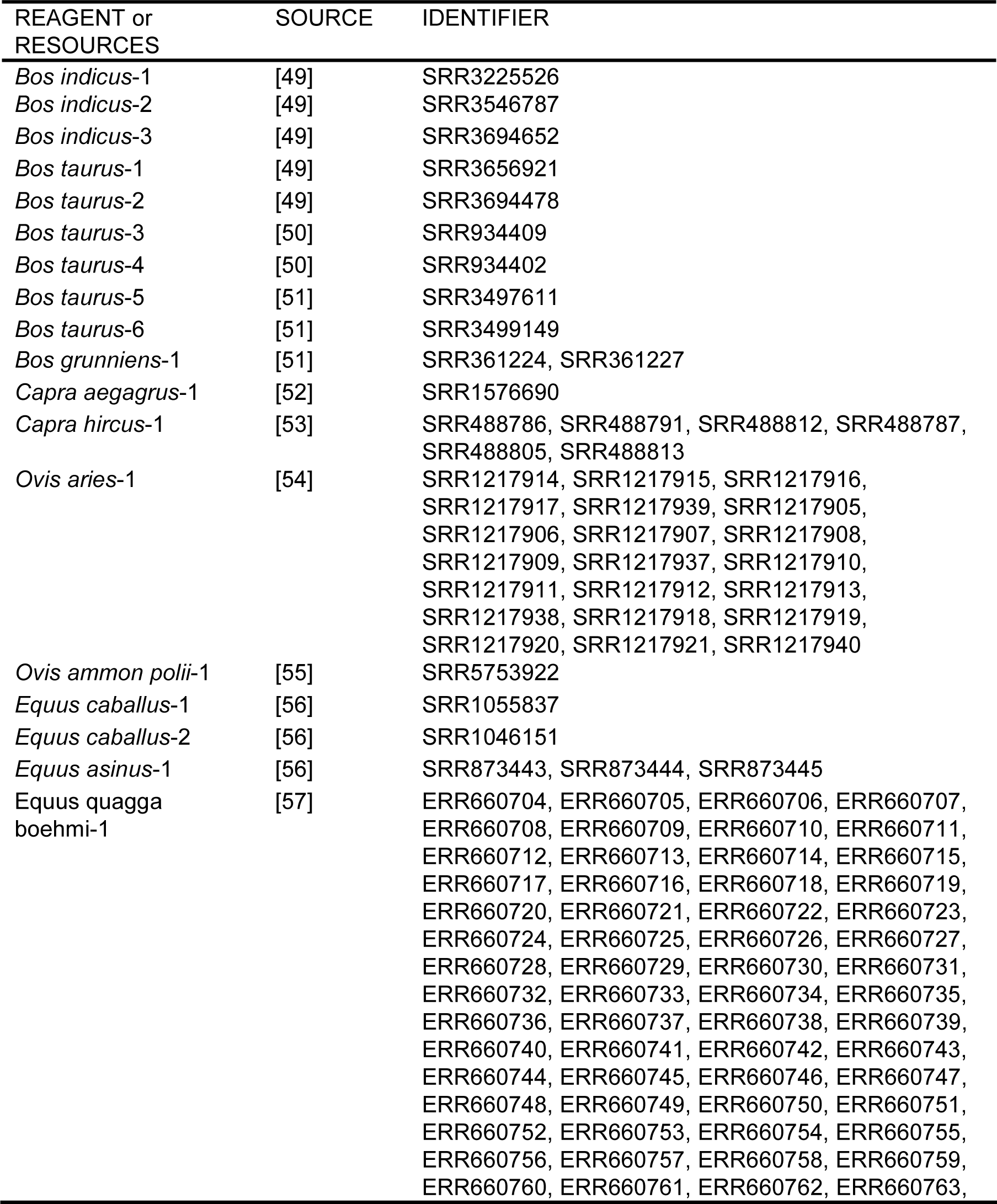
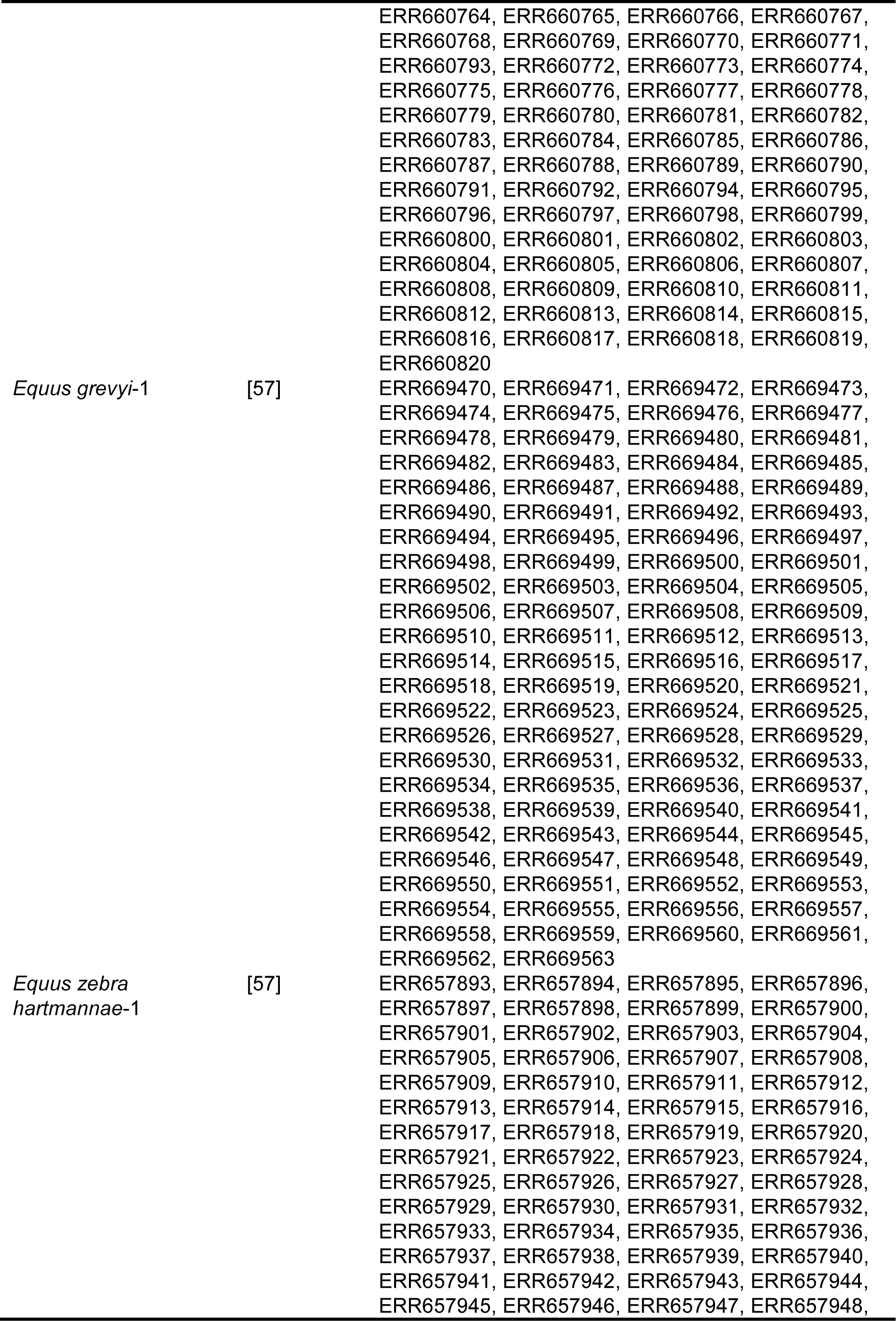
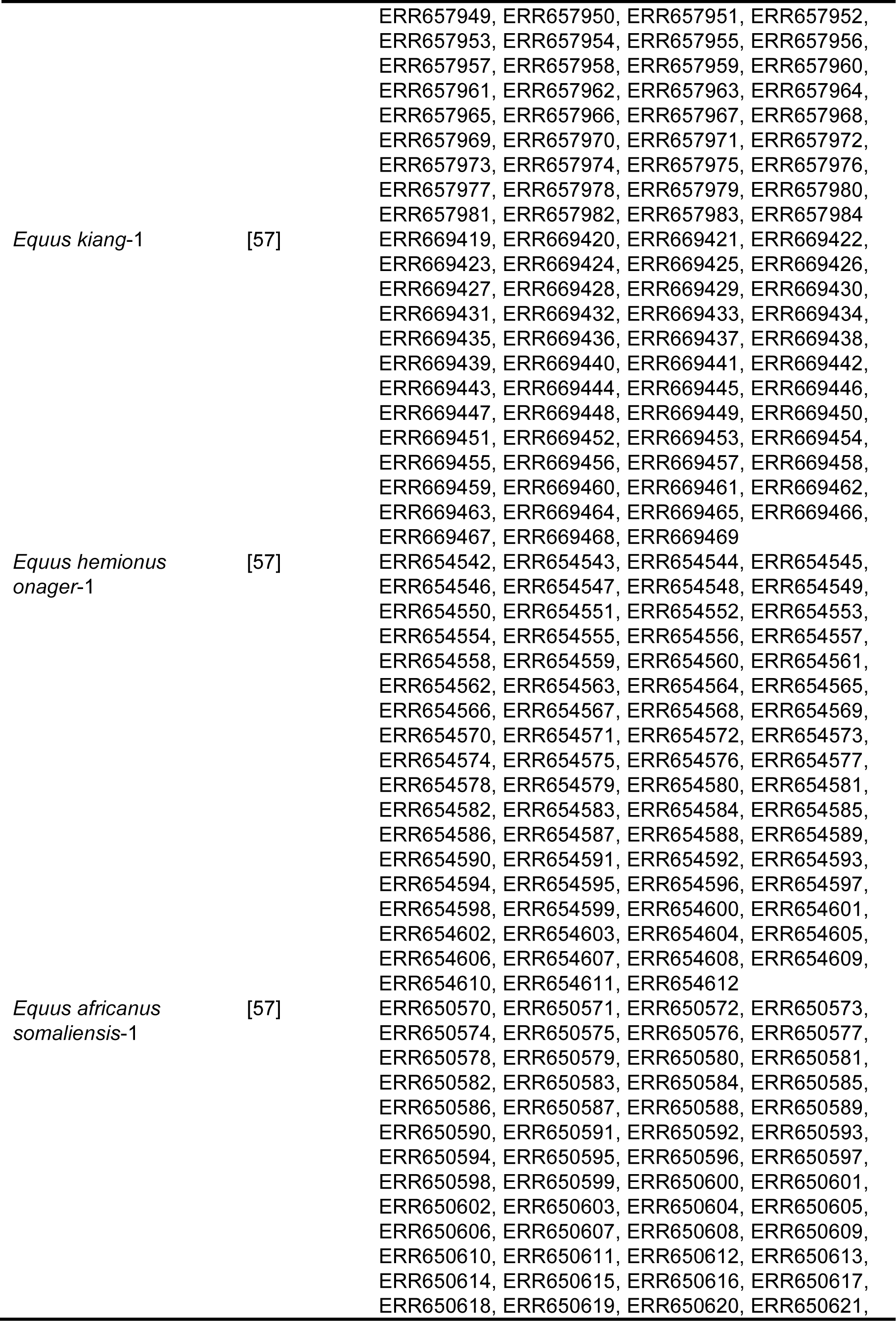
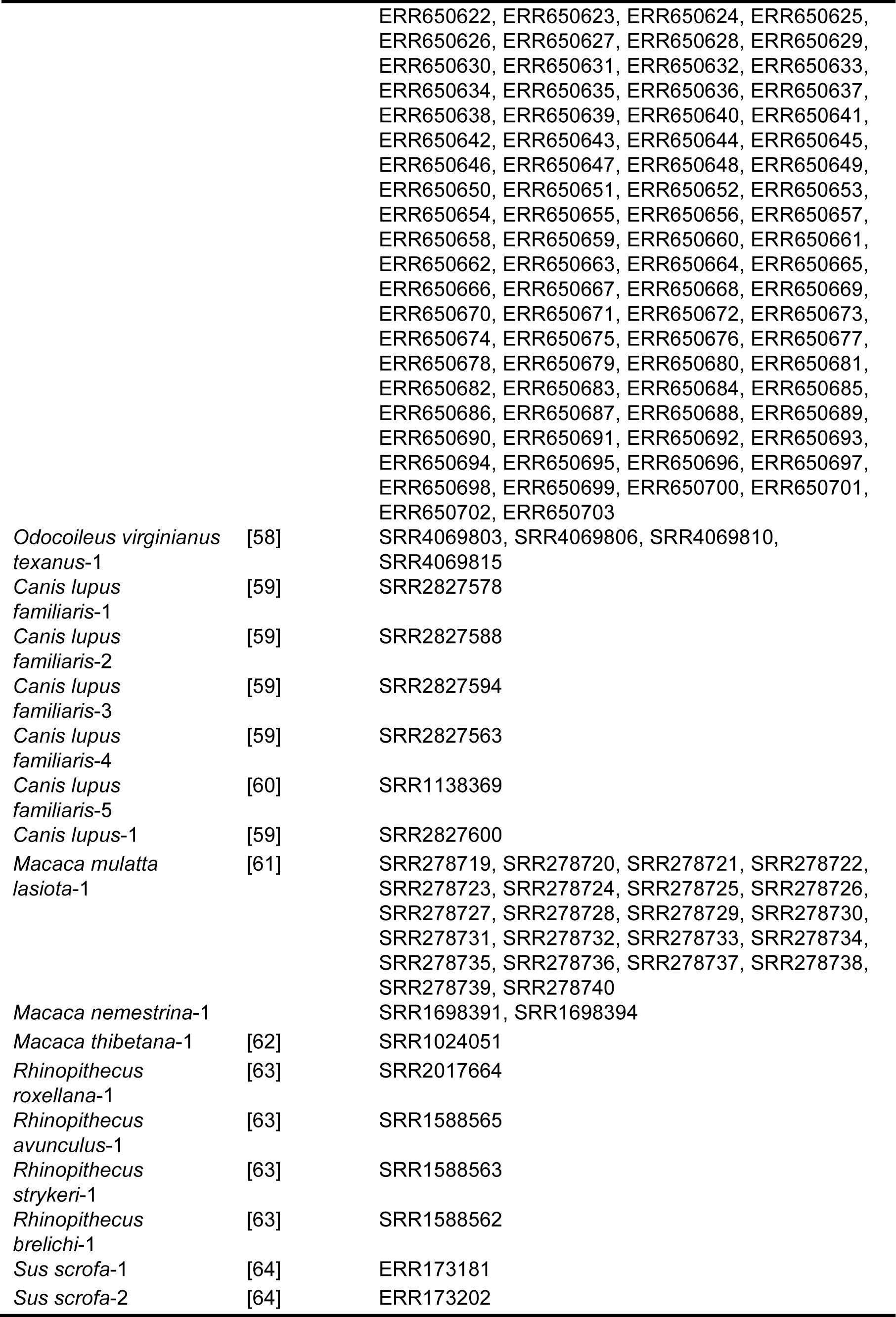
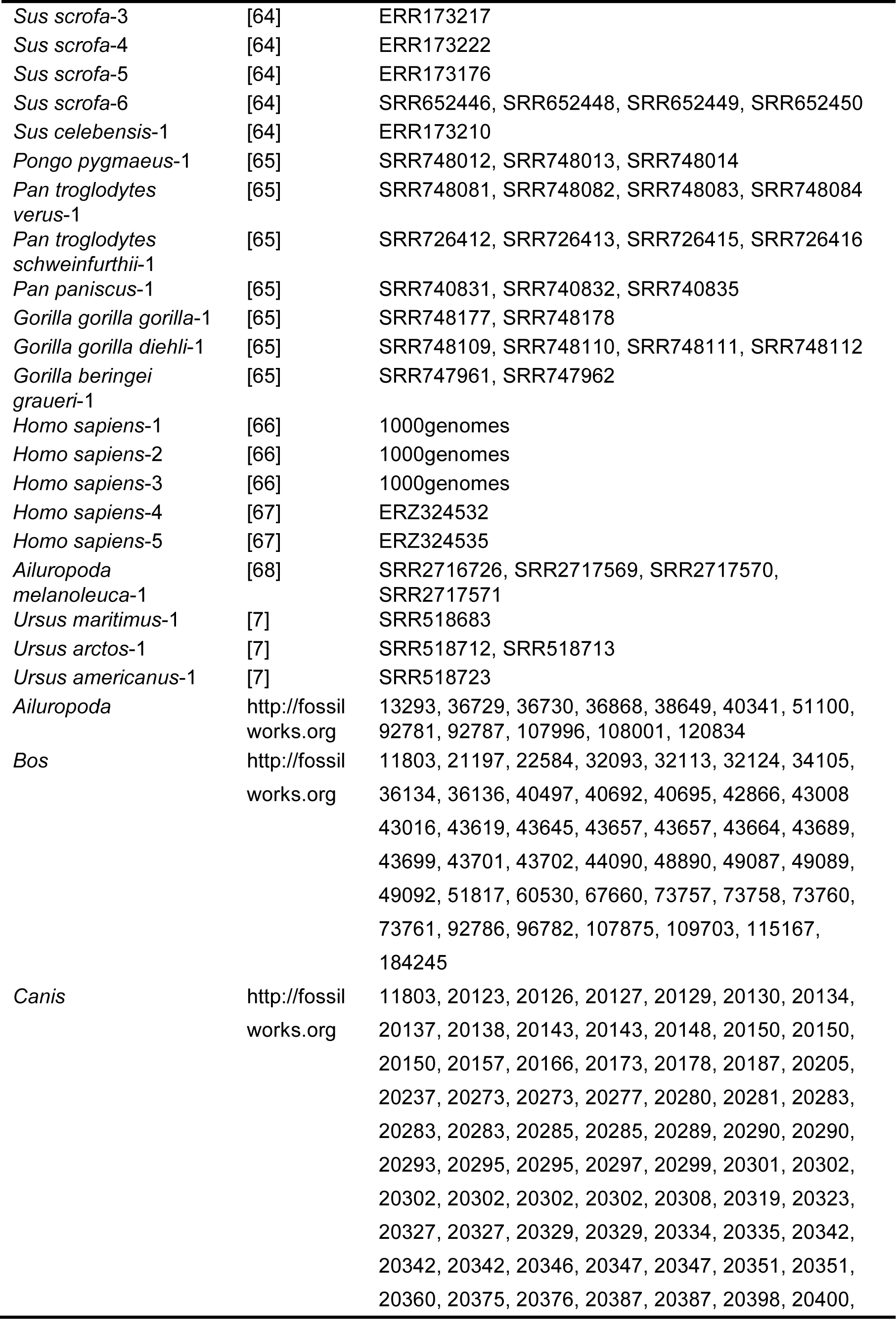
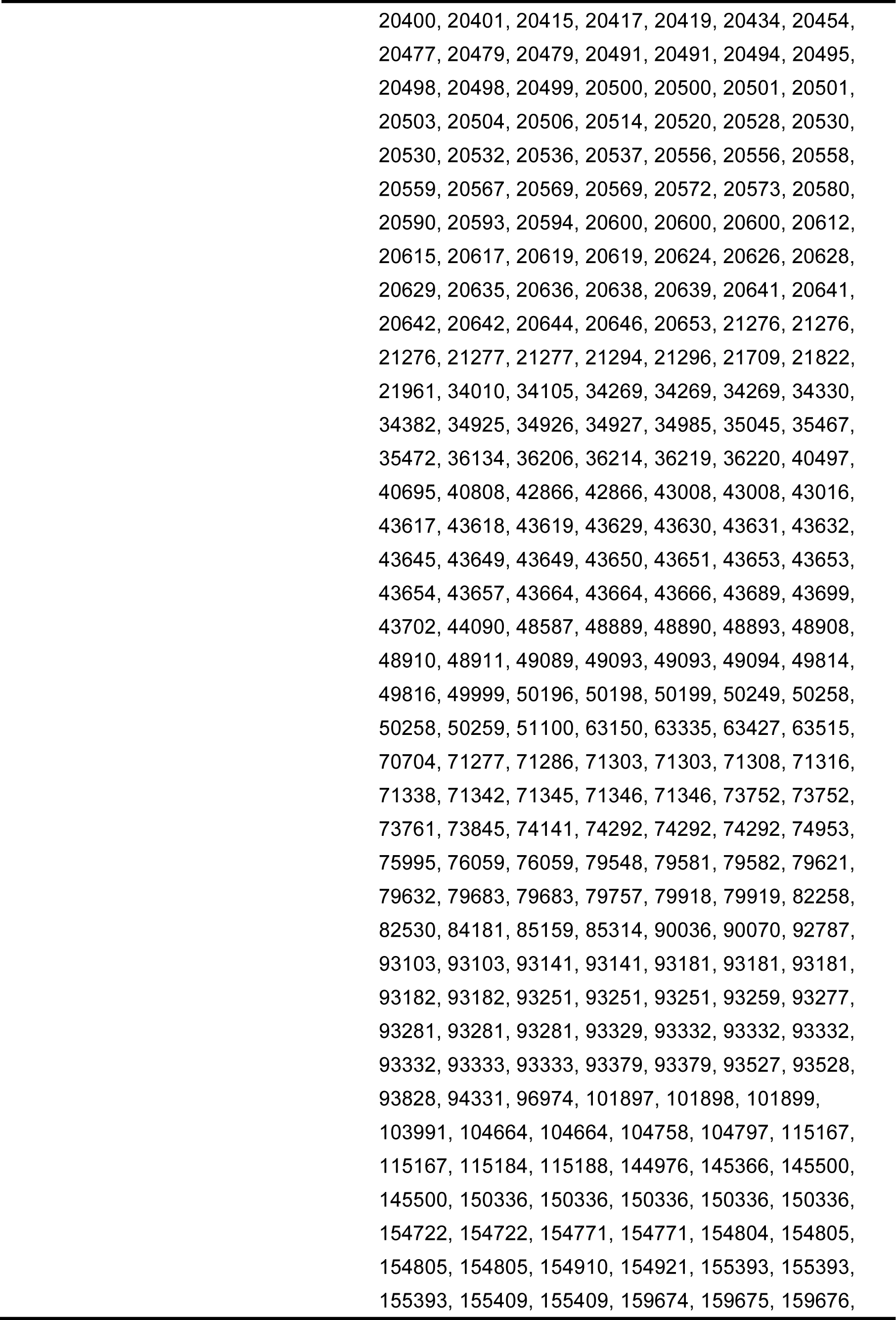
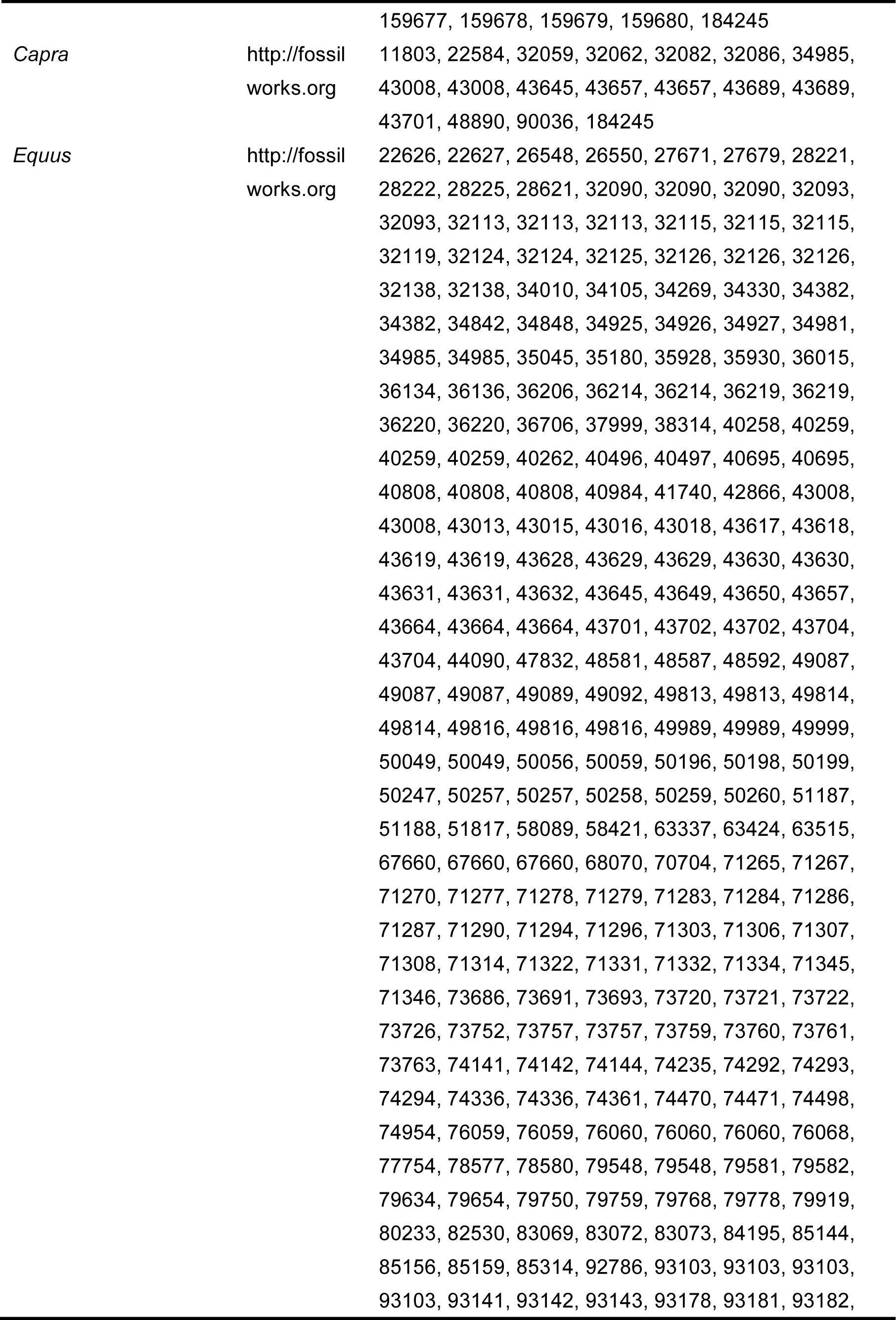
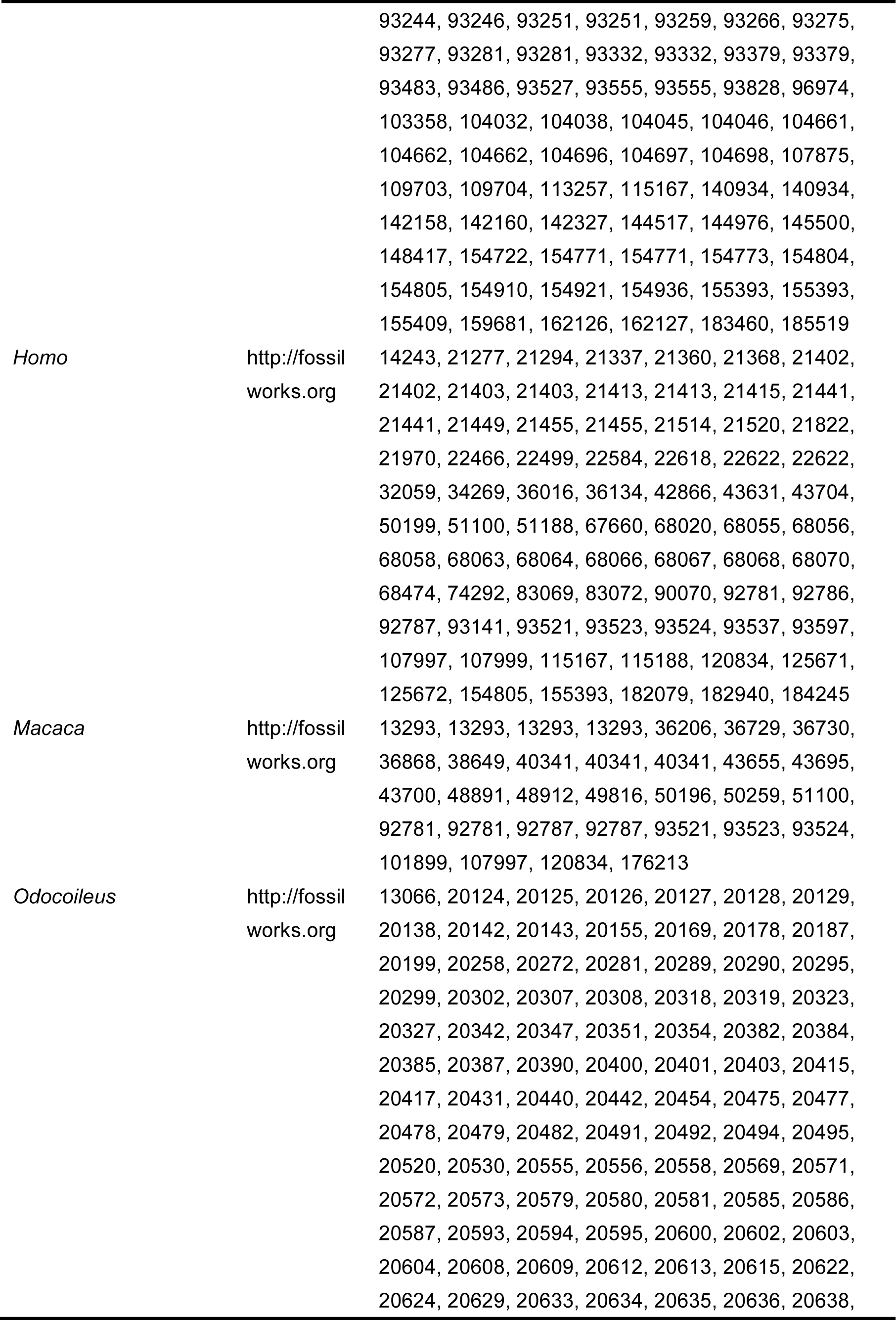
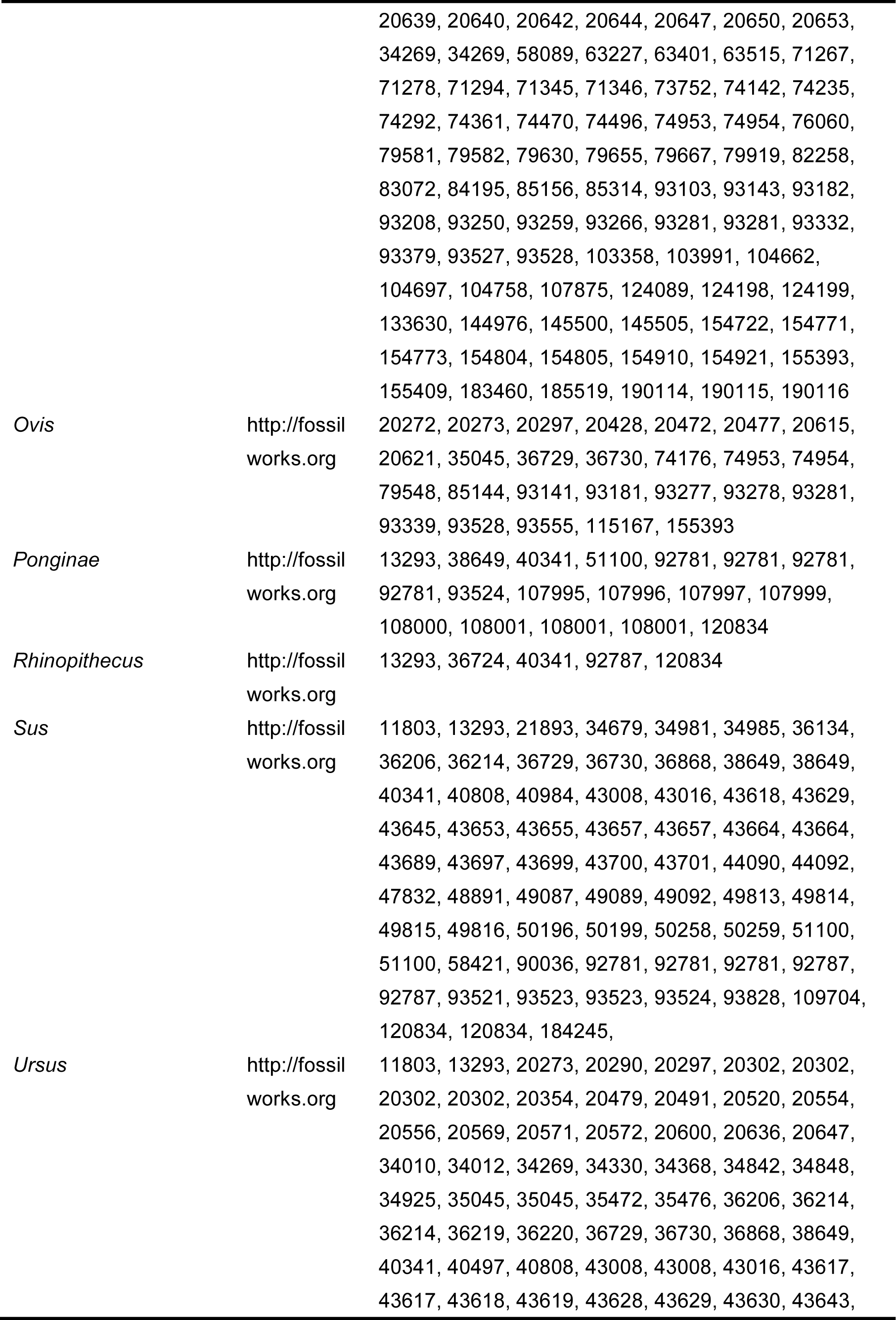
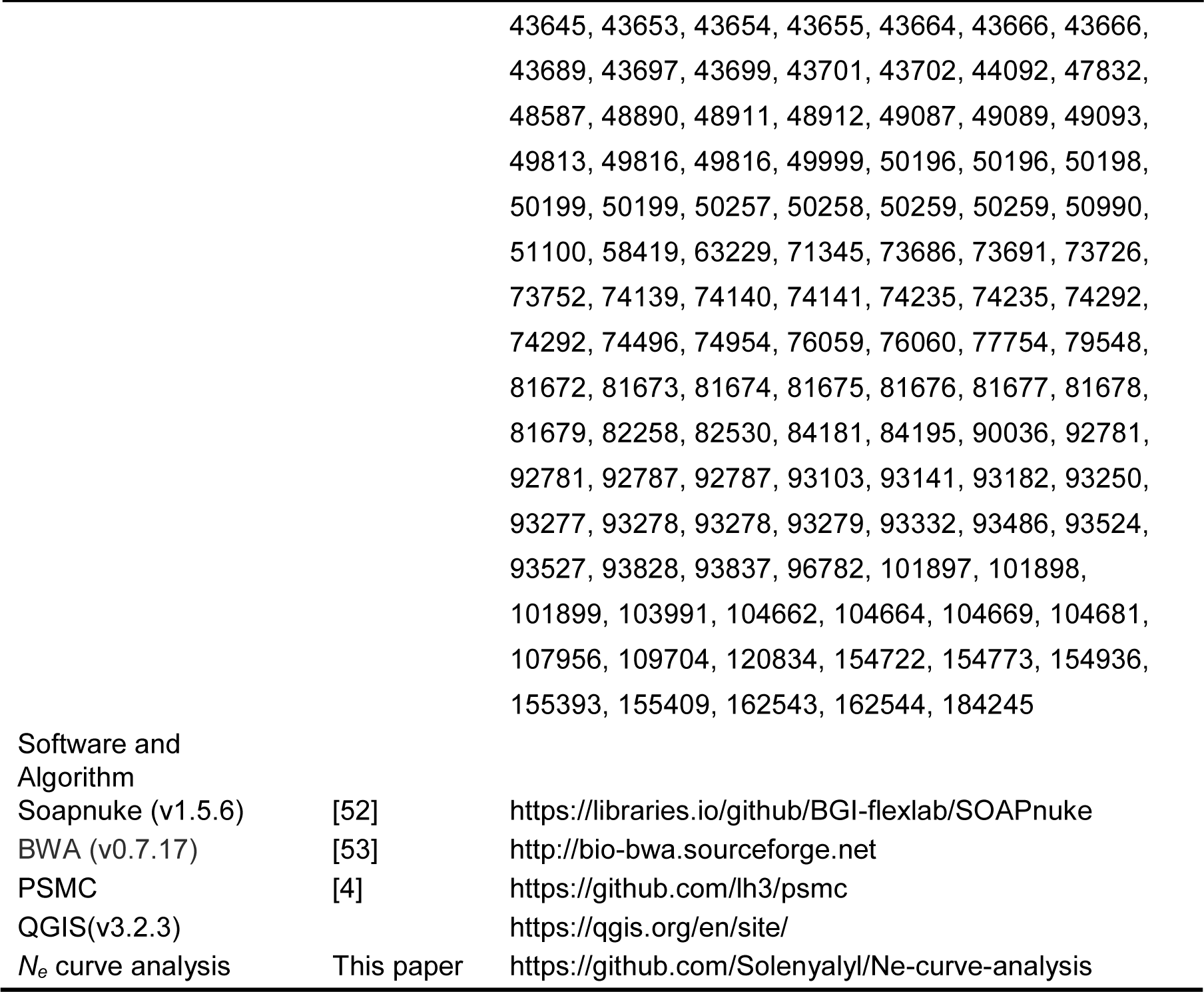

## CONTACT FOR REAGENT AND RESOURCE SHARING

Further information and requests for resources and reagents should be directed to and will be fulfilled by the Lead Contact, Shuaicheng Li.

## METHOD DETAILS

### Mammal and fossil distribution

Mammalian SRA data came from different studies, and not all of their locations were recorded. Mammalian place of origin was regarded as the location if a sample was collected from the zoo; domestication location was taken to be the location if it was assured; and when it was ambiguous, sampling location was regarded as the location. For each of the remaining samples without location recorded, we researched the species distribution in the world and then chose a representative place as the location. Our mammals were distributed among 15 genera, and their fossil records were found in http://fossilworks.org/ except for *Pan* and *Gorilla*. Those fossils originated in the Calabrian (1.8-0.781 mya), middle Pleistocene (0.781-0.126 mya) or late Pleistocene (0.126-0.0117 mya). Mammal location and fossil data distribution were plotted using QGIS (Figure S1 and Figure S3).

### Data collection and preprocessing

We downloaded mammalian Illumina sequence data from public SRA datasets (https://www.ncbi.nlm.nih.gov/sra/), Chinese BAM files were downloaded from https://www.ebi.ac.uk/, and non-Chinese BAM files were collected from http://www.internationalgenome.org/. Including 60 specimens from 35 species, these SRA data were sequenced by Illumina paired-end sequencing technology using different sequencing platforms, therefore our first step was transforming all the Illumina sequence data into uniform Q+33 Illumina format. All the raw sequencing reads were filtered by Soapnuke [52] (http://soap.genomics.org.cn/) and each was aligned to its respective reference genome or the most related genome from the same genus by BWA 0.7.17 [53]. Samples with genome coverage depth less than 6X were excluded.

### N_e_ curve generation

The historical *N*_*e*_ curve of mammals was inferred by PSMC based on genome data [4]. A PSMC-based consensus genome sequence was obtained by using ‘mpileup’ in SAMTOOLS. When the RMS (root-mean-squared) quality of reads was below 20, their covered sites were marked N for missing data. The sites whose read depths were less than a third or more than twice the average depth across the genome were masked to avoid collapsed regions in the assembly. When we ran PSMC, the upper limit of TRMCA was set to 15, the initial θ/ρ value was set to 5 and time interval was set to ‘1*4+25*2+1*4+1*6’ as Li and Durbin [4] did. The mutation rate and generation time we applied are in Table S3. The genome coverage depth, coverage ratio, and false-negative ratio are in Table S2. For statistical analysis of these *N*_*e*_ curves see sections below.

## QUANTIFICATION AND STATISTICAL ANALYSIS

### PC analysis and hierarchical clustering

PCA was used to identify the principal components of *N*_*e*_ curves. Every *N*_*e*_ curve contains 58 points. We standardized these data to unit standard deviation and centralized these points approximately 0, then extracted their principal components to conduct hierarchical clustering. The distance matrix between samples was calculated from Euclidean distance and distance between two clusters was calculated by the UPGMA (unweighted pair group method with arithmetic mean) method.

### N_e_ curve extrema analysis

To obtain the extrema of every curve, we marked increasing trends as ‘1’ and decreasing trends as ‘-1’ in curves, then the latter trend number was used to subtract the former trend number to obtain the extrema. ‘-2’ and ‘2’ represent crest and trough, respectively. To verify whether these extrema are distributed randomly, we compared these extrema with a uniform distribution to obtain Hopkins statistics. After obtaining a clustering trend, these extrema were clustered by the K-means method to get a pattern. Then we used this pattern to predict every curve extrema into seven clusters by removing those neglectable extrema (difference within 0.1).

## DATA AND SOFTWARE AVAILABILITY

N_e_ curve analysis code is available from https://github.com/Solenyalyl/Ne-curve-analysis.

**Table S1. *N*_*e*_ curve extrema seven clusters (related to Figure 2,3)**

**Table S2. coverage depth, coverage ratio, and false-negative ratio of 60 mammal genomes(related to Figure 1-4)**

