## supplementary for "Pleistocene Mammal Population Fluctuation Patterns Inferred by Their Genomes"

| Sample name | Mutation rate (per generation per site) | Generation time(year) | Reference |
|---|---|---|---|
| *Bos indicus*-1 | 9.8x10-9 | 5 | [S1] |
| *Bos indicus*-2 | 9.8x10-9 | 5 | [S1] |
| *Bos indicus*-3 | 9.8x10-9 | 5 | [S1] |
| *Bos taurus*-1 | 9.8x10-9 | 5 | [S1] |
| *Bos taurus*-2 | 9.8x10-9 | 5 | [S1] |
| *Bos taurus*-3 | 9.8x10-9 | 5 | [S1] |
| *Bos taurus*-4 | 9.8x10-9 | 5 | [S1] |
| *Bos taurus*-5 | 9.8x10-9 | 5 | [S1] |
| *Bos taurus*-6 | 9.8x10-9 | 5 | [S1] |
| *Bos grunniens*-1 | 9.8x10-9 | 5 | [S1] |
| *Capra aegagrus*-1 | 3.89x10-9 | 2 | [S2] |
| *Capra hircus*-1 | 3.89x10-9 | 2 | [S2] |
| *Ovis aries*-1 | 3.89x10-9 | 2 | [S2] |
| *Ovis ammon polii*-1 | 3.89x10-9 | 2 | [S2] |
| *Equus caballus*-1 | 8x10-9 | 5 | [S3] |
| *Equus caballus*-2 | 8x10-9 | 5 | [S3] |
| *Equus asinus*-1 | 8x10-9 | 5 | [S3] |
| *Equus quagga boehmi*-1 | 8x10-9 | 5 | [S3] |
| *Equus grevyi*-1 | 8x10-9 | 5 | [S3] |
| *Equus zebra hartmannae*-1 | 8x10-9 | 5 | [S3] |
| *Equus kiang*-1 | 8x10-9 | 5 | [S3] |
| *Equus hemionus onager*-1 | 8x10-9 | 5 | [S3] |
| *Equus africanus somaliensis*-1 | 8x10-9 | 5 | [S3] |
| *Odocoileus virginianus texanus*-1 | 6.6x10-9 | 3 | [S4] |
| *Canis lupus familiaris*-1 | 6.6x10-9 | 3 | [S5] |
| *Canis lupus familiaris*-2 | 6.6x10-9 | 3 | [S5] |
| *Canis lupus familiaris*-3 | 6.6x10-9 | 3 | [S5] |
| *Canis lupus familiaris*-4 | 6.6x10-9 | 3 | [S5] |
| *Canis lupus familiaris*-5 | 6.6x10-9 | 3 | [S5] |
| *Canis lupus*-1 | 6.6x10-9 | 3 | [S5] |
| *Macaca mulatta lasiota*-1 | 2.5x10-8 | 6 | [S6, S7] |
| *Macaca nemestrina*-1 | 2.5x10-8 | 6 | [S6, S7] |
| *Macaca thibetana*-1 | 2.5x10-8 | 6 | [S6, S7] |
| *Rhinopithecus roxellana*-1 | 5x10-9 | 5 | [S8] |
| *Rhinopithecus avunculus*-1 | 5x10-9 | 5 | [S8] |
| *Rhinopithecus strykeri*-1 | 5x10-9 | 5 | [S8] |
| *Rhinopithecus brelichi*-1 | 5x10-9 | 5 | [S8] |
| *Sus scrofa*-1 | 1.2x10-8 | 5 | [S9] |
| *Sus scrofa*-2 | 1.2x10-8 | 5 | [S9] |
| *Sus scrofa*-3 | 1.2x10-8 | 5 | [S9] |
| *Sus scrofa*-4 | 1.2x10-8 | 5 | [S9] |
| *Sus scrofa*-5 | 1.2x10-8 | 5 | [S9] |
| *Sus scrofa*-6 | 1.2x10-8 | 5 | [S9] |

| | | | |
|---|---|---|---|
| *Sus celebensis*-1 | 1.2x10-8 | 5 | [S9] |
| *Pongo pygmaeus*-1 | 1.2x10-8 | 26 | [S5, S10] |
| *Pan troglodytes verus*-1 | 1.2x10-8 | 25 | [S5, S10] |
| *Pan troglodytes schweinfurthii*-1 | 1.2x10-8 | 25 | [S5, S10] |
| *Pan paniscus*-1 | 1.2x10-8 | 25 | [S5, S10] |
| *Gorilla gorilla gorilla*-1 | 1.2x10-8 | 19 | [S5, S10] |
| *Gorilla gorilla diehli*-1 | 1.2x10-8 | 19 | [S5, S10] |
| *Gorilla beringei graueri*-1 | 1.2x10-8 | 19 | [S5, S10] |
| *Homo sapiens*-1 | 1.2x10-8 | 25 | [S5, S11] |
| *Homo sapiens*-2 | 1.2x10-8 | 25 | [S5, S11] |
| *Homo sapiens*-3 | 1.2x10-8 | 25 | [S5, S11] |
| *Homo sapiens*-5 | 1.2x10-8 | 25 | [S5, S11] |
| *Homo sapiens*-4 | 1.2x10-8 | 25 | [S5, S11] |
| *Ailuropoda melanoleuca*-1 | 1.29×10-8 | 12 | [S12] |
| *Ursus maritimus*-1 | 1×10-8 | 10 | [S13] |
| *Ursus arctos*-1 | 1×10-8 | 10 | [S13] |
| *Ursus americanus*-1 | 1×10-8 | 10 | [S13] |

**Table S4. mammalian mutation rate and generation time (related to Figure 1-4).** Those informations were gathered from the references.

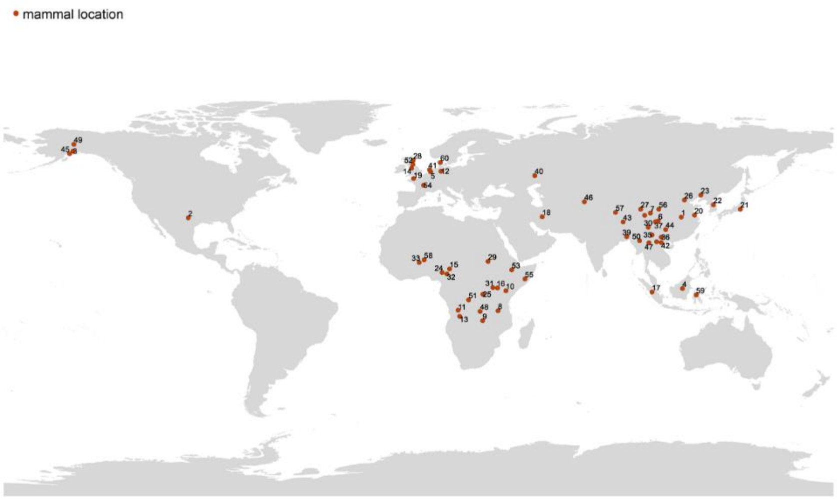


**Figure S1. mammal distributions (related to Figure 1-4).** The red point represents location, the correspondence of sample and name shows below:

| sample | number |
|---|---|
| *Homo sapiens*-5 | 1 |
| *Odocoileus virginianus texanus*-1 | 2 |
| *Ursus americanus*-1 | 3 |
| *Pongo pygmaeus*-1 | 4 |
| *Bos taurus*-3 | 5 |
| *Ailuropoda melanoleuca*-1 | 6 |
| *Equus hemionus onager*-1 | 7 |

| | |
|---|---|
| *Equus asinus*-1 | 8 |
| *Equus quagga boehmi*-1 | 9 |
| *Equus grevyi*-1 | 10 |
| *Pan troglodytes schweinfurthii*-1 | 11 |
| *Equus caballus*-2 | 12 |
| *Equus zebra hartmannae*-1 | 13 |
| *Homo sapiens*-2 | 14 |
| *Bos taurus*-2 | 15 |
| *Bos indicus*-2 | 16 |
| *Sus scrofa*-5 | 17 |
| *Capra aegagrus*-1 | 18 |
| *Bos taurus*-5 | 19 |
| *Sus scrofa*-2 | 20 |
| *Homo sapiens*-1 | 21 |
| *Bos taurus*-4 | 22 |
| *Canis lupus*-1 | 23 |
| *Canis lupus familiaris*-3 | 24 |
| *Gorilla beringei graueri*-1 | 25 |
| *Sus scrofa*-4 | 26 |
| *Bos grunniens*-1 | 27 |
| *Bos taurus*-6 | 28 |
| *Bos indicus*-3 | 29 |
| *Canis lupus familiaris*-5 | 30 |
| *Bos taurus*-1 | 31 |
| *Gorilla gorilla diehli*-1 | 32 |
| *Homo sapiens*-3 | 33 |
| *Capra hircus*-1 | 34 |
| *Canis lupus familiaris*-2 | 35 |
| *Macaca mulatta lasiota*-1 | 36 |
| *Macaca thibetana*-1 | 37 |
| *Rhinopithecus roxellana*-1 | 38 |
| *Macaca nemestrina*-1 | 39 |
| *Ursus maritimus*-1 | 40 |
| *Ovis aries*-1 | 41 |
| *Rhinopithecus avunculus*-1 | 42 |
| *Equus kiang*-1 | 43 |
| *Rhinopithecus brelichi*-1 | 44 |
| *Ursus arctos*-1 | 45 |
| *Ovis ammon polii*-1 | 46 |
| *Homo sapiens*-4 | 47 |
| *Gorilla gorilla gorilla*-1 | 48 |
| *Canis lupus familiaris*-4 | 49 |
| *Rhinopithecus strykeri*-1 | 50 |
| *Pan paniscus*-1 | 51 |

| | |
|---|---|
| *Equus caballus*-1 | 52 |
| *Bos indicus*-1 | 53 |
| *Sus scrofa*-3 | 54 |
| *Equus africanus somaliensis*-1 | 55 |
| *Canis lupus familiaris*-1 | 56 |
| *Sus scrofa*-6 | 57 |
| *Pan troglodytes verus*-1 | 58 |
| *Sus celebensis*-1 | 59 |
| *Sus scrofa*-1 | 60 |

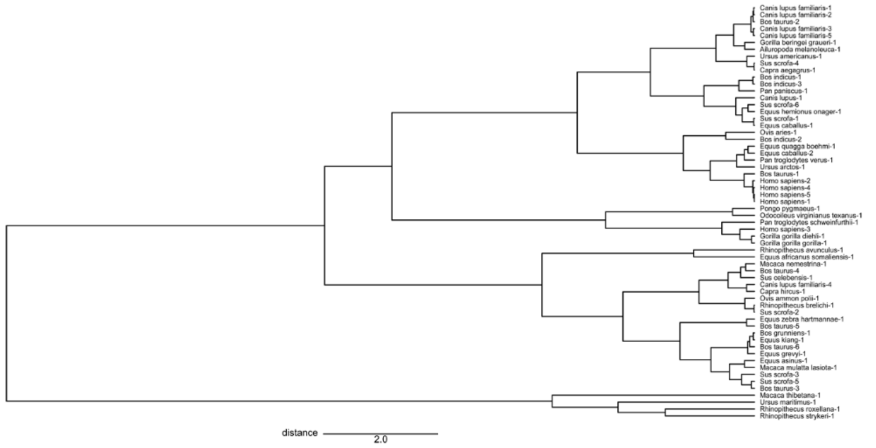


**Figure S2. Hierarchical cluster of $N_e$ curve PC1 (related to Figure 1).**

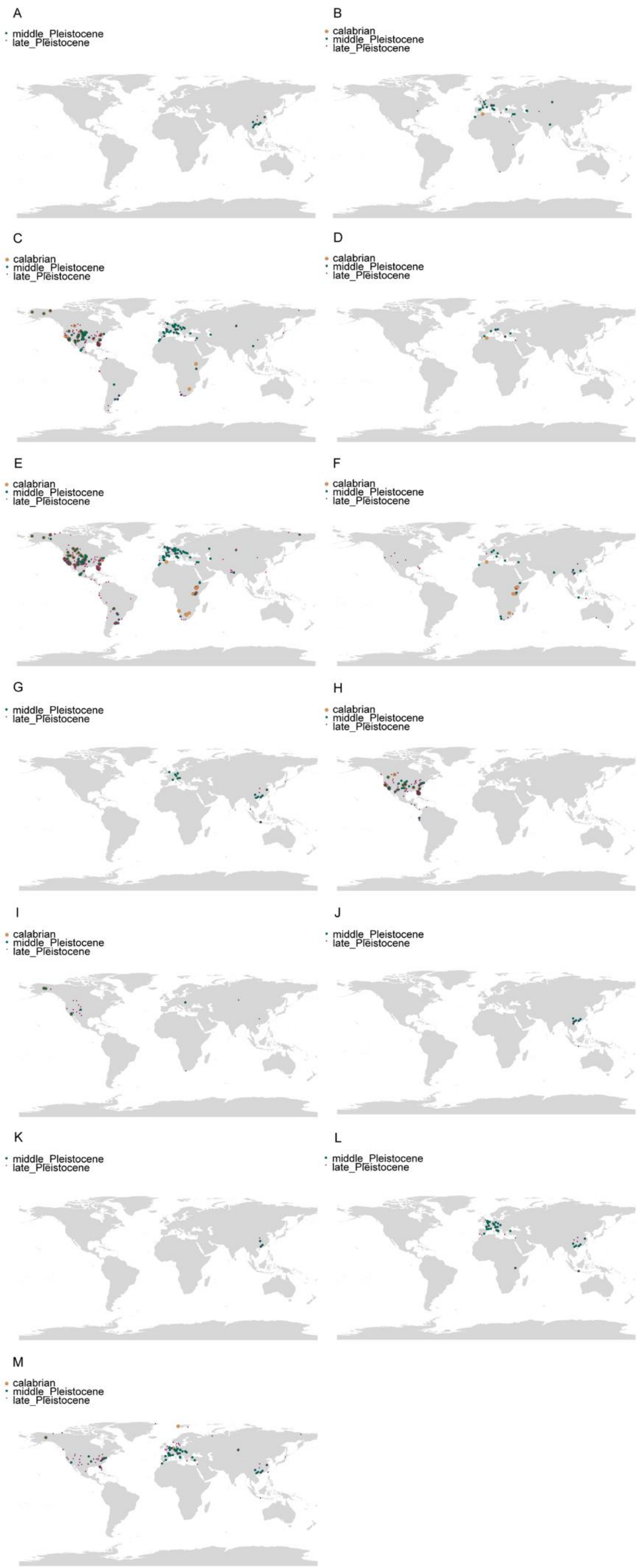
A
middle_Pleistocene
late_Pleistocene
B
calabrian
middle_Pleistocene
late_Pleistocene
C
calabrian
middle_Pleistocene
late_Pleistocene
D
calabrian
middle_Pleistocene
late_Pleistocene
E
calabrian
middle_Pleistocene
late_Pleistocene
F
calabrian
middle_Pleistocene
late_Pleistocene
G
middle_Pleistocene
late_Pleistocene
H
calabrian
middle_Pleistocene
late_Pleistocene
I
calabrian
middle_Pleistocene
late_Pleistocene
J
middle_Pleistocene
late_Pleistocene
K
middle_Pleistocene
late_Pleistocene
L
middle_Pleistocene
late_Pleistocene
M
calabrian
middle_Pleistocene
late_Pleistocene

**Figure S3. Mammalian fossil record distributions across the world in the Pleistocene (related to Figure 2-3).** The scatters represent fossil record locations. Big orange circle, middle green circle and small pink circle represent fossils came from the Calabrian, the MP and the LP respectively. (A) *Ailuropoda*. (B) *Bos*. (C) *Canis*. (D) *Capra*. (E) *Equus*. (F) *Homo*. (G) *Macaca*. (H) *Odocoileus*. (I) *Ovis*. (J) *Ponginae*. (K) *Rhinopithecus*. (L) *Sus*. (M) *Ursus*.

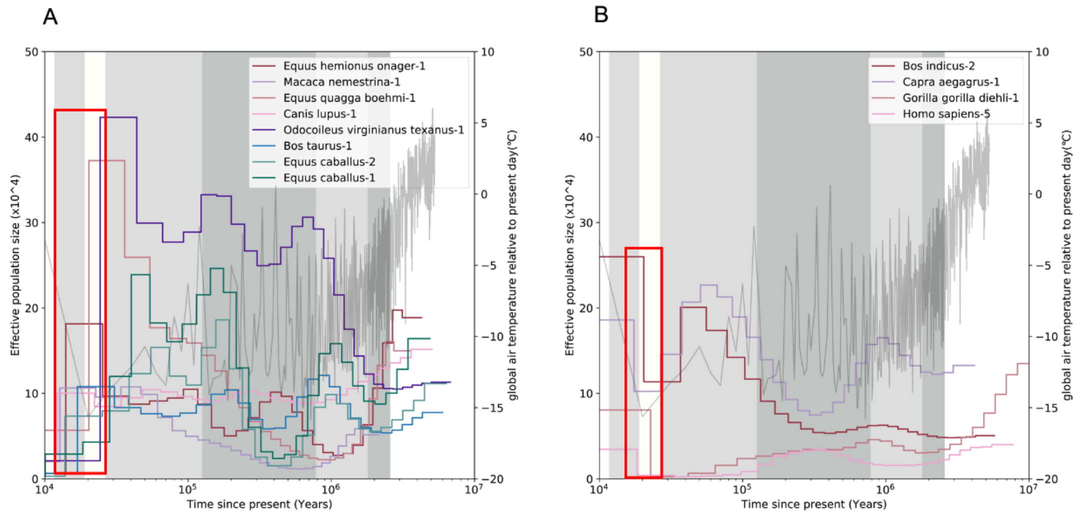


**Figure S4. Mammal $N_e$ curves with sharp fluctuation during the Late Pleistocene (related to Figure 4).** (A) PSMC results show a sharp decline for eight mammals around the LGM; (B) PSMC results show a sharp increase for 4 mammals around the LGM. Every curve represents a mammal effective population size fluctuation. The snow-white region indicates the LGM and gray or light gray regions indicate 4 stages in the Pleistocene. The dark gray line in the background represents temperature fluctuation in the Pleistocene. In the red frame, the mammal population displays an upheaval.
